# Foraging behavior across paths that vary in risk cues and frequency of occurrence

**DOI:** 10.1101/2019.12.17.878819

**Authors:** Emily K. Lessig, Peter Nonacs

## Abstract

Cooperatively foraging species often adjust their search strategies in complex environments to efficiently find and exploit food sources. These strategies become more complicated when food and risk can be simultaneously present and when they differ in predictability. For example, there may be multiple paths to reach a foraging site that vary in risk. This study examines how colonies of Argentine ants (*Linepithema humile*) respond to such a situation where identical-length paths differ in how they present risk. The risk cues are either a live competitor (velvety tree ants, *Liometopum occidentale* (LO)) or formic acid (FA), a defensive chemical commonly associated with formicine ant species. Across four paths to food, the presence of cues also varied from always to never present. Although the non-risky path was used more often, in no case did colonies completely avoid the paths with risk. Overall, more *L. humile* workers explored paths associated with LO than with FA. This had a significant impact on foraging ability where LO colonies were faster at finding food than FA colonies. Further, L. *humile* workers’ response to FA was similar over time while declined for LO, suggesting a ‘dear enemy’ habituation and reduction in aggression over time. Thus, it appears that *L. humile* foragers categorize risk cues and will vary their responses in potentially effective ways.

## Introduction

Efficient food acquisition challenges all animals. Foragers navigate complex environments that present a variety of costs and benefits, often associated both with the distribution or quality of food patches and with the travel associated to reach them. Patch exploitation and movements between patches can be influenced by travel distance, presence of competing species or cooperating conspecifics, food quality, and hunger levels (Alatalo and Lundberg 1986; Anderson 1984; Beckers et al. 1990; Conradt and Roper 2005; Vittori et al. 2006; Ronconi and Burger 2011; Yates and Nonacs 2016; Yamada 2017). Given that a forager’s first goal is to effectively encounter potential food items, the time it takes to discover food has implications for foraging success and efficiency (Beverly 2009). In heterogeneous environments, organisms may need to change their search strategies in order to efficiently find food. For example, Argentine ants *(Linepithema humile)* appear to prioritize rapid recruitment to food once it is found, rather than maximizing food encounter rates when search areas differ in spatial complexity and food appears ephemerally (Denton and Nonacs, 2018).

Foraging strategies and spatial distributions of workers can also vary in response to the risk that is present along a path. *Lasius pallatarsis* ant colonies abandon patches with associated mortality risk to forage at “safer” patches (Nonacs and Dill, 1988). In *Formica* ants (*F. perpilosa and F. integroides*), smaller foragers avoid sites at which risk is present and larger foragers may spend more time at these sites in a defensive posture (Kay and Rissing, 2005; Tanner, 2008). Maximizing foraging effort therefore requires recognizing key properties of different patches and allocating foraging efforts across patches in a manner that increases time spent at “good” patches and decreases time spent at “bad.” In heterogenous or changing landscapes, animals may then need to sample their environment and develop expectations about future encounters to efficiently exploit the area (Stephens and Krebs 1986). This process of combining older information with newer information to alter expectations is known as Bayesian updating (Valone, 2006) and has been observed in a wide variety of taxonomic groups (e.g. Lima 1984, 1985; Valone & Brown 1989; Valone 1991, 1992; Krebs & Inman 1992; Alonso et al. 1995; Olsson et al. 1999; van Gils et al. 2003; Stamps et al. 2018).

Cooperatively foraging species, such as ants, add two more aspects to foraging strategies. First, individual workers are ‘disposable’ in the sense that their deaths are expected and do not directly influence the reproduction of the colony. Such foragers gain indirect fitness by helping kin reproduce and may behave very differently from animals that risk their own reproductive success (Nonacs and Dill, 1990). Second, many ant species mark and maintain trails that can denote the optimal route to a food source; balancing trade-offs between various qualities amongst trails (Nonacs and Dill, 1990,1991, Latty et al. 2017).

In this study, we examine the path choice that ants make over time when: (1) There are multiple paths of equal length to a food source, and (2) negative stimuli (i.e., cues that a competing species may be nearby) are present along these paths in different frequencies, ranging from 0% (never present) to 100% (always present). Previous studies have focused on decisions between food and risk that are presented simultaneously, but here we focus on how colonies learn and respond over time to negative features of their environment that may vary in how often and predictably they are present.

## Methods

### Collection

We setup six replicate colonies of *L. humile* containing approximately 500-800 workers, 6-8 queens, and brood collected at the University of California, Los Angeles. The ants nested in open plastic containers, filled with molded plaster of Paris, which was kept moist for nest humidity, and coated on the sides with Fluon. Water was provided *ad libitum*.

### Experiment

The experimental arenas consisted of four equidistant paths made of clear, plastic tubing to a single foraging arena (distance = 90cm). To measure the time it took ants to find food, we added sugar water to the foraging arena daily between approximately 12:00 and 15:00 h. Time was recorded as how long it took the first ant to find the food and was recorded up to the first 10 minutes. Food was removed two hours after placement (whether or not ants were foraging) to avoid colonies becoming satiated. To present the ants with negative stimuli, we placed cells at the midway point of each path where cues were added (Fig. 1). Workers from three colonies each encountered either several workers of the aggressive velvety tree ant (*Liometopum occidentale* (LO)) (Hoey-Chamberlain et al. 2013) or formic acid (FA) on the paths and depending on the treatment. LO and FA were placed along paths between 8:00 and 11:00 h. daily. They were present along paths for approximately 24 hours and not correlated with the appearance of food. The four paths differed such that LO or FA were present 0, 25, 50 or 100% of trials. Therefore, to reach food the *L. humile* foragers always had one path that never had risk associated, another that always had risk, and two that might or might not have had risk associated with them. This means that for any given trial, 1-3 paths could have had LO or FA present.

**Figure 1.**
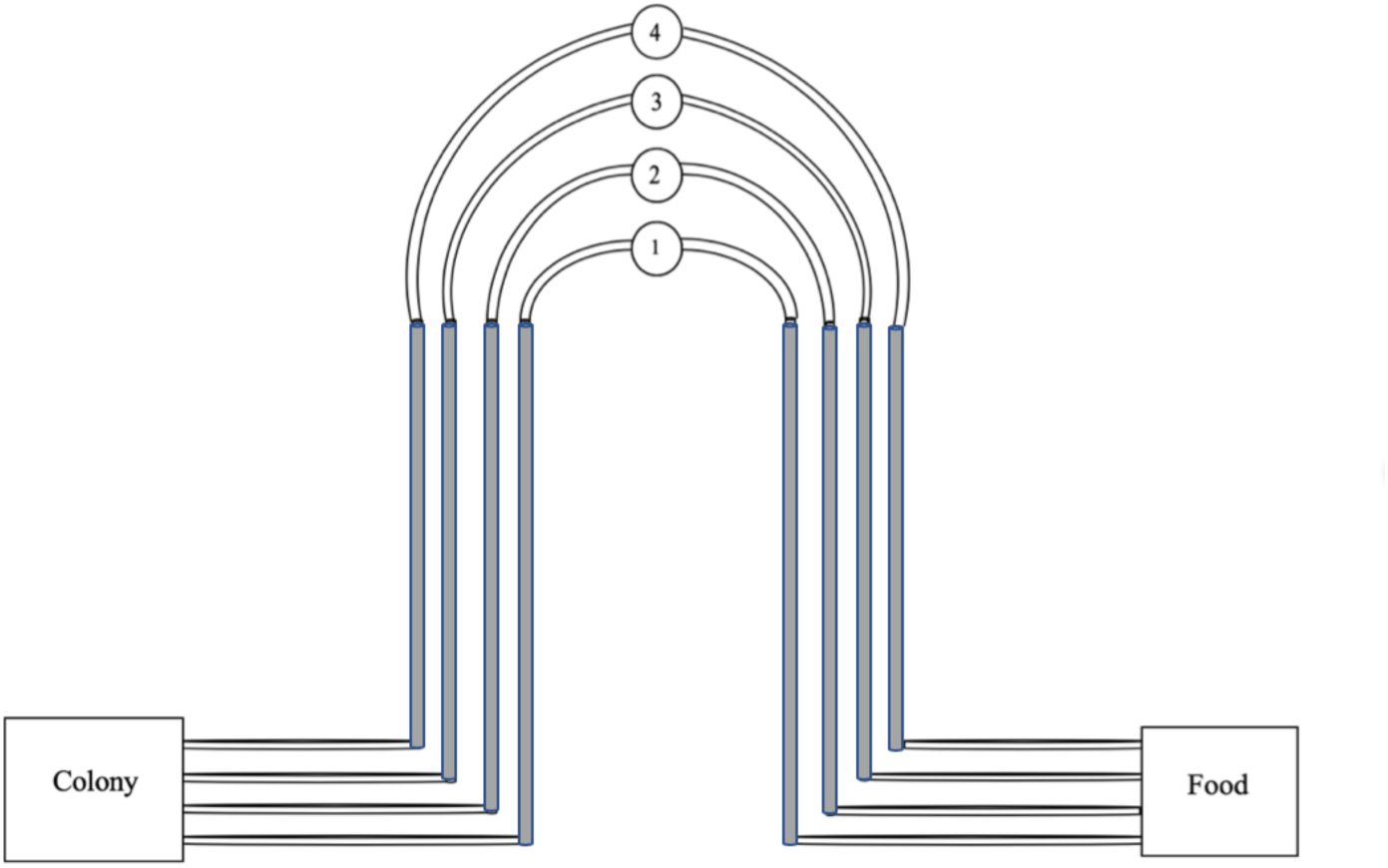
Diagram of the experimental grid. Cells labeled 1-4 indicated where negative stimuli were placed. All paths were of equal length.

We collected LO from Descanso Gardens (Los Angeles, CA) and housed them in a similar manner to that of *L. humile*. Both *L. humile* and *L. occidentale* are in the same subfamily of Dolichoderinae. FA is the primary defensive chemical used by ant species in the subfamily Formicinae (Hefetz & Blum 1978; Blum 1978). Colonies of ants were counted every four to five days to ensure numbers did not drop drastically and workers were added if numbers dropped below 500. Trials lasted approximately 20 days to allow each colony to experience all frequencies of negative stimuli at least four times.

#### Risk cue: LO

We placed approximately 15-20 LO workers in cells 1-4 (Fig. 1). Mesh was present between the LO workers and the *L. humile* workers, to prevent physical contact (Fig. S2). However, they were able to exchange chemical cues. LO workers were removed and replaced daily to ensure their numbers and effect were consistent over the course of the experiment.

#### Risk cue: FA

We placed cotton pads with 100 µL of FA (Walmart, USA) in cells 1-4 (Fig. 1). We covered the cells with lids for these trials to prevent the FA from completely dissipating over time but cut holes (diameter =2.54cm) into the encounter chambers for aeration and to prevent FA fumes from accumulating in deadly amounts. We removed pads and/or replaced them daily depending on the presentation schedule. If FA was to be absent for the day, we cleaned cells to remove any leftover traces.

### Video Observations

We programmed webcams to take fifteen-minute recordings of path use, approximately three times per day: one at least three hours prior to food presentation (morning observation), one while food was present (afternoon observation), and one in the evening, at least three hours after food had been removed (evening observation). The number of observations across colonies varied because they were not all set up simultaneously and some observations had to be discarded due to ants escaping or the clarity of video taken. Observations were at minimum 3 hours apart and at approximately the same time daily. We later scored these videos to examine path use over time.

### Video Scoring

Only the number of ants crossing the shaded portion of the experimental grids (Fig. 1) was recorded as this was most detectable across all cameras and colonies. To avoid double-counting an ant on a path, we kept spatial track of a forager’s location and did not count it more than once when simply moving back and forth on the same path.

## Statistics

We conducted linear-mixed effect models to study the amount of time ants took to find food across days. Stimulus type was included in the model as well, as *L. humile* workers could respond differently to LO or FA affecting finding times. We began by generating a null model containing colony as a random intercept, then iteratively incorporated fixed effects into the model, in order to find which combination of fixed effects generated the lowest AIC value for the model. We conducted an ANOVA to examine mean time to find food across days by stimulus type (FA or LO), all in R version 3.5.2 (R Core Team 2018).

We also conducted an ANOVA to examine path use of ants per day averaged across the number of observations per day (log-transformed) against risk cue (LO or FA), path traveled (0, 25, 50 or 100), number of paths with risk cues on a given day (i.e. “danger,” ranging from 1-3), and day of the experiment. Not every colony was tested on every day and therefore for statistical analysis, time was divided into quartiles of 6 days each.

## Results

In the analysis examining time to find food across days, including stimulus type as a fixed effect significantly improved model fit (Table 1). This indicates that type of stimulus predicts the amount of time it takes ants to find food. Further, in the analysis examining mean time to find food across days by stimulus type, food was found significantly faster in LO treatments than in FA treatments (Fig. 2; p < 0.0001). This indicates that foraging behavior (i.e. time to find food) is influenced by stimulus type.

**Table 1:**
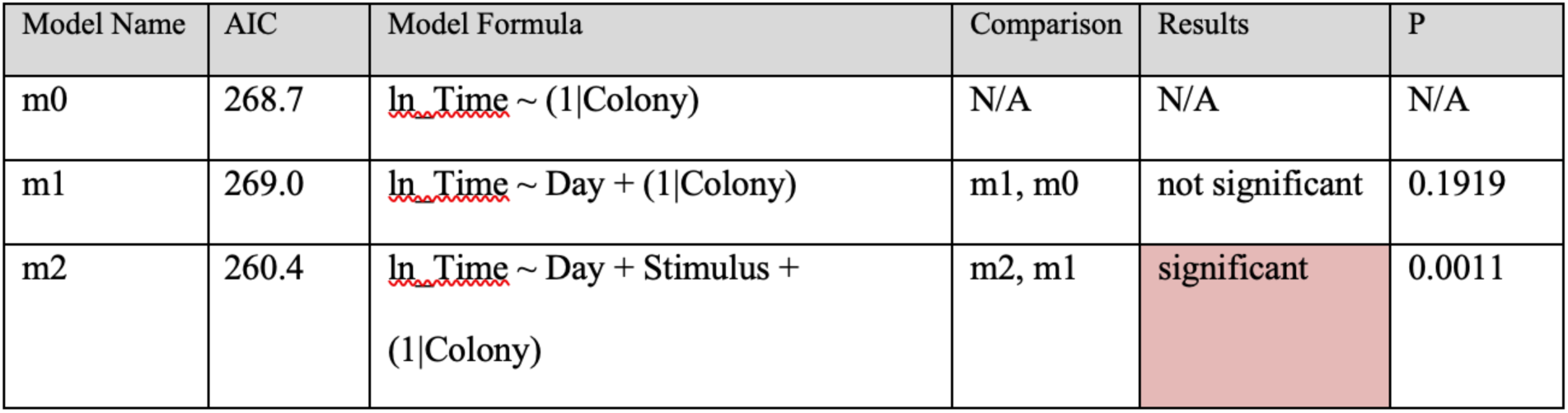
Model comparison for time data across days. Model m2 was chosen based on AIC value and comparison with other models using likelihood ratio tests.

**Figure 2:**
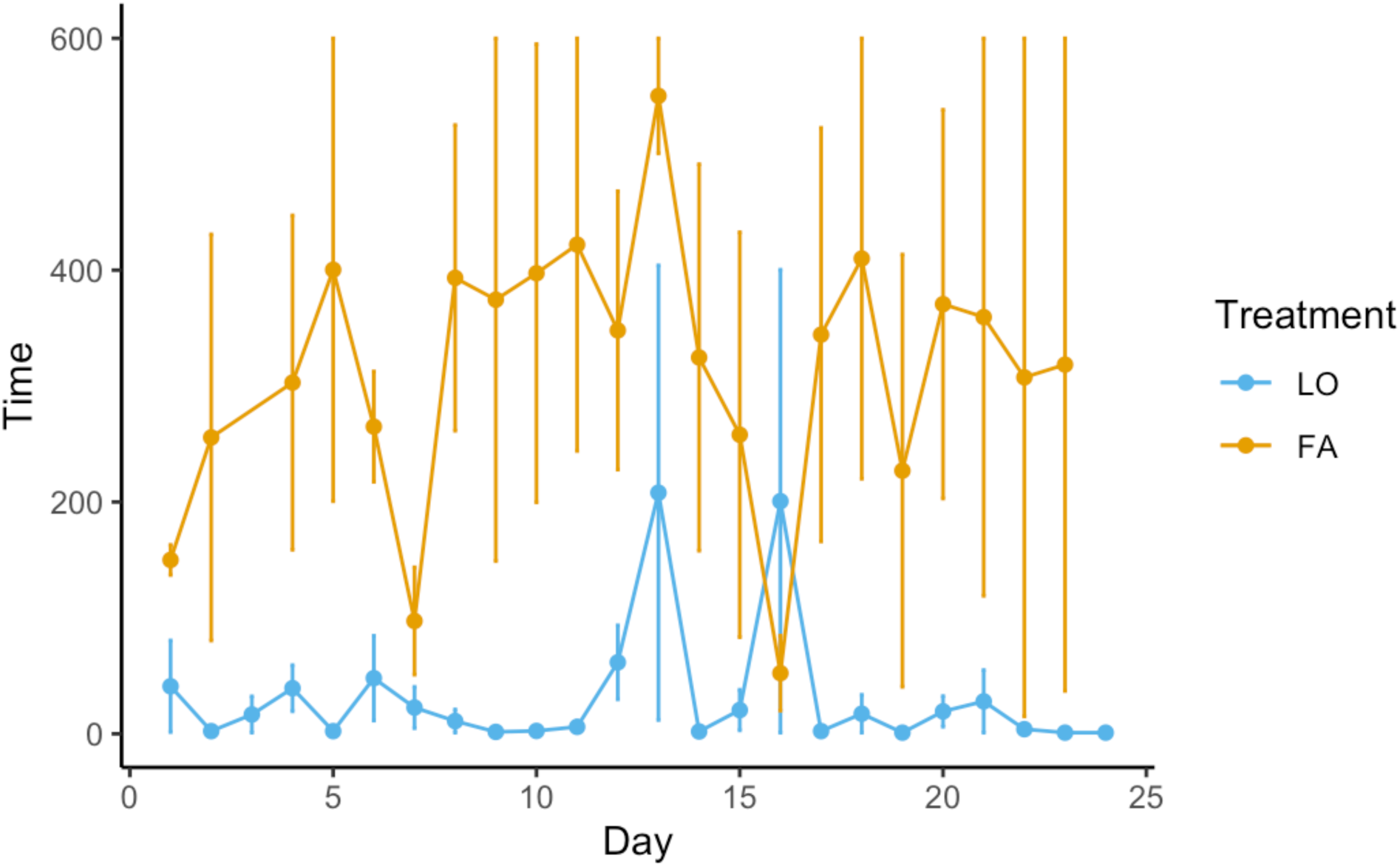
Mean time to find food across days for all four paths by stimulus type (where colony 1-3 received FA and 4-6 received LO). There was a significant difference in time to find food between colonies that received FA as opposed to colonies that received LO (p < 0.0001).

In the analysis examining number of ants per day against risk cue, path traveled, danger (in terms of number of paths with risk cues present on a given day), and day of the experiment, danger had neither a significant main effect nor any significant interaction effects. Therefore, it was dropped as a factor in the ANOVA. In regard to the other main effects, significantly more ants were on the paths with LO as opposed to FA (Fig. 3; p < 0.0001, F = 154.87, df = 1). The frequency of risk appearing on the paths also significantly affected their use (Fig. 3; p < 0.0004, F = 6.205, df = 3), where there was a significant difference between the number of ants using the 0 and 25% paths (p < 0.0061); the 0 and 50% paths (p < 0.0001); and the 50 and 100% paths (p < 0.0061). Additionally, the day of the experiment had a significant affect, where the overall number of ants declined over time (Fig. 3; p < 0.0001; F = 9.681; df = 3).

**Figure 3:**
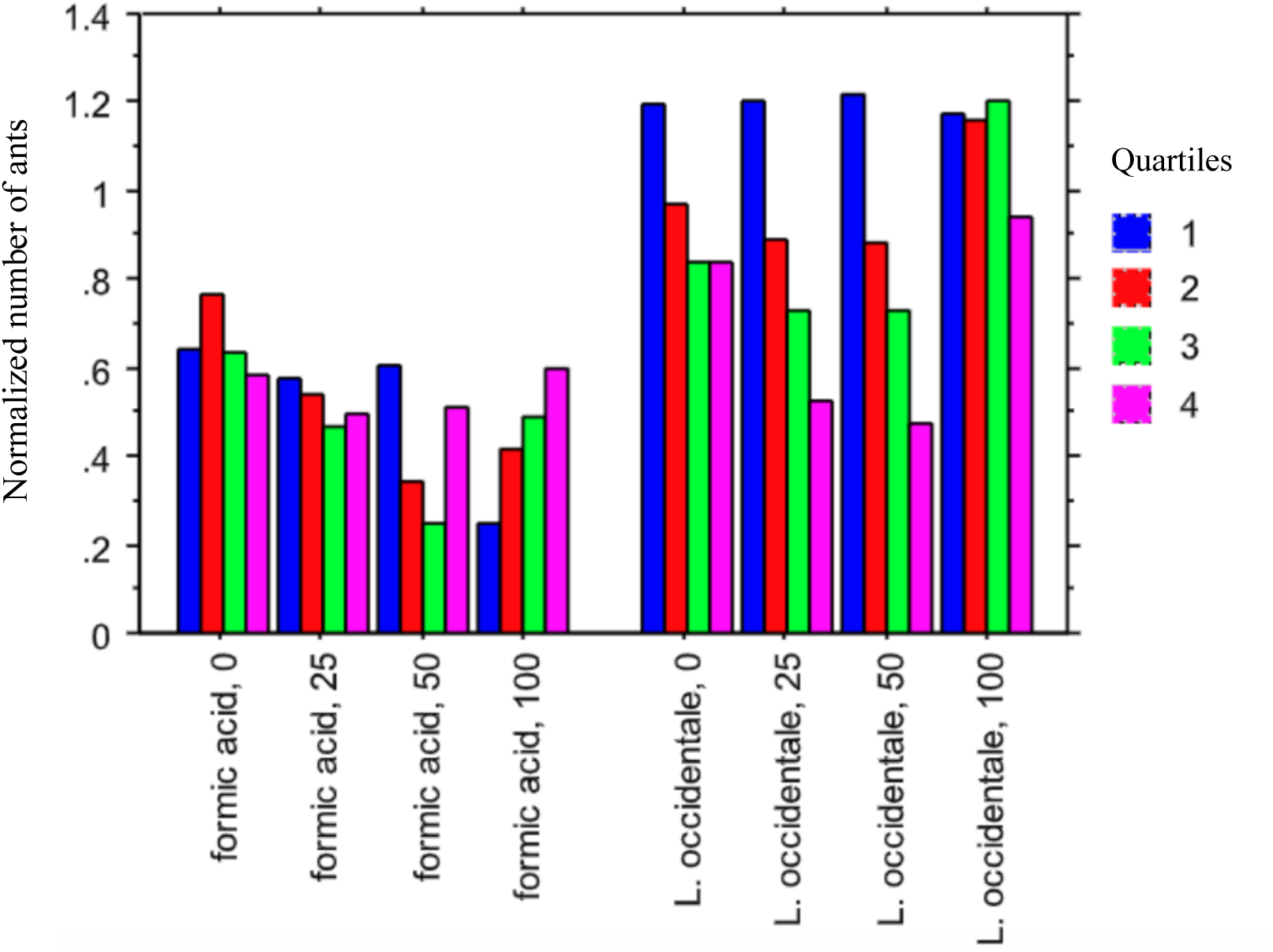
Mean number of ants per day averaged across the number of observations per day (log-transformed) against risk cue (LO or FA), path traveled (0, 25, 50 or 100), number of paths with risk cues on a given day (i.e. “danger,” ranging from 1-3), and day of the experiment (divided into quartiles). Colors respond to the division of days into 4 quartiles.

When examining the 2-way interactions, we found when FA is the stimulus the number of ants declined as risk got more frequent. However, for LO, the consistently riskiest path (100%) drew the most ants (Fig. 3; p < 0.0002, F = 6.639, df = 3). Further, the FA response was similar over time, while the LO response consistently declined (Fig. 3; p < 0.0001, F= 10.130, df = 3). Except for the 100% path, the number of ants declined over time (Fig. 3: p < 0.0026; F = 2.872, df = 9). The 3-way interaction between stimulus, path, and time was not significant.

## Discussion

Cooperatively foraging species are a model system to examine how individuals with limited knowledge and cognitive capabilities can achieve complex goals such as navigating complex environments. Patch exploitation and movements between patches can be influenced by a variety of factors including presence of competing species. While a forager’s goal is to successfully find food items, it also has to balance safety along paths. In heterogenous environments, organisms may need to change their search strategies in order to efficiently find food as well stay safe. Previous studies have focused on decisions between food and risk that are presented simultaneously, but here we focus on how colonies learn and respond to negative features of their environment that may vary in how often and how predictably they are present.

Different types of risk can also elicit different responses. In our experiments, we found the two types of cues about risk drew significantly different responses both in regard to foraging behavior and path use. Ants patrol/defend areas more intensely when in the presence of live LO workers than with only FA, resulting in more ants on paths with LO as compared to FA. We can attribute the lower numbers on paths with FA due to the effectiveness of this chemical weapon.

FA as a defense in formicine species is more effective in conflicts with Argentine ants than the defenses of the dolichoderine velvety tree ants. This has been noted in *Nylanderia fulva* and other formicines where formic acid as a chemical weapon achieves competitive dominance in combat with *Solenopsis invicta* (LeBrun et al. 2014). Thus, even the occasional presence of FA may depress Argentine ant activity in that area or path. As FA is a particularly effective chemical weapon in combat, learning which trails to use or avoid FA is an important aspect of their foraging strategy.

Further, the large numerical difference in response along paths with live LO workers as compared to FA likely resulted in LO colonies finding food when it appeared significantly faster than FA colonies. This suggests that food discovery rate by Argentine ants can be directly affected by their responses to encountering different competitive species. Additionally, their response to FA is similar over time while for LO, their response declines over time. This suggests that the lack of direct contact and fights results in habituation and a ‘dear enemy’ reduction in aggression over time (Langen et al. 2000) as *L. humile* workers decline in their response to LO over time.

Additionally, the predictability of the risk along paths also had a significant effect on path use. The always safe (0%) path is used more overall than the paths with risk. Further, the overall number of ants on the paths declines over time, except for the never safe (100%) path. This suggests that ants can assess and respond to risk present along paths and they moderate their responses to paths that are not consistently risky but maintain a more consistent presence when in response to paths that are always risky.

This work demonstrates that Argentine ants are able to learn about their environment and use this information to effectively navigate and efficiently exploit their environment. Further, this work elucidates how individual ants (with local knowledge and the ability to interact) can scale up to effective organizations that optimally achieve vital, but potentially competing, objectives. This is due to the fact that learning at the colony level is often more effective than at the individual level as workers can vary in age and experience, and colonies allow for communication and an increase in numbers. These results are relevant to understanding how groups coordinate and function in a wide variety of species.

