## Supplementary Figures for "Foraging behavior across paths that vary in risk cues and frequency of occurrence"

**Supplementary Information**

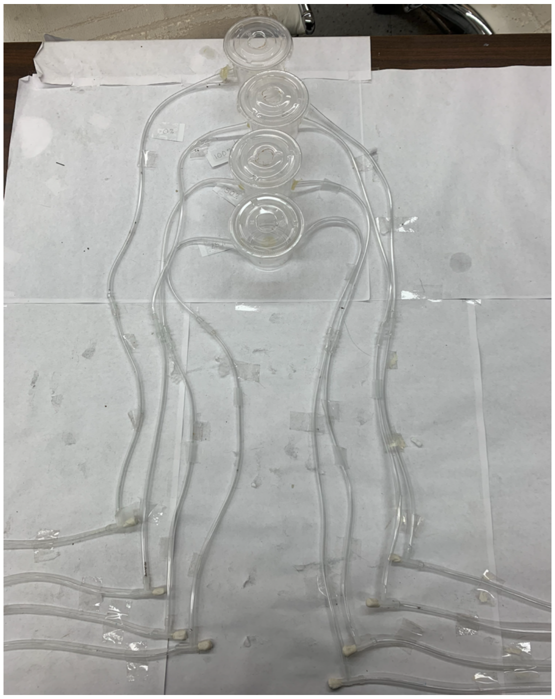

**Fig. S1:** Picture of experimental setup, including paths that were scored and cells where stimuli were placed.

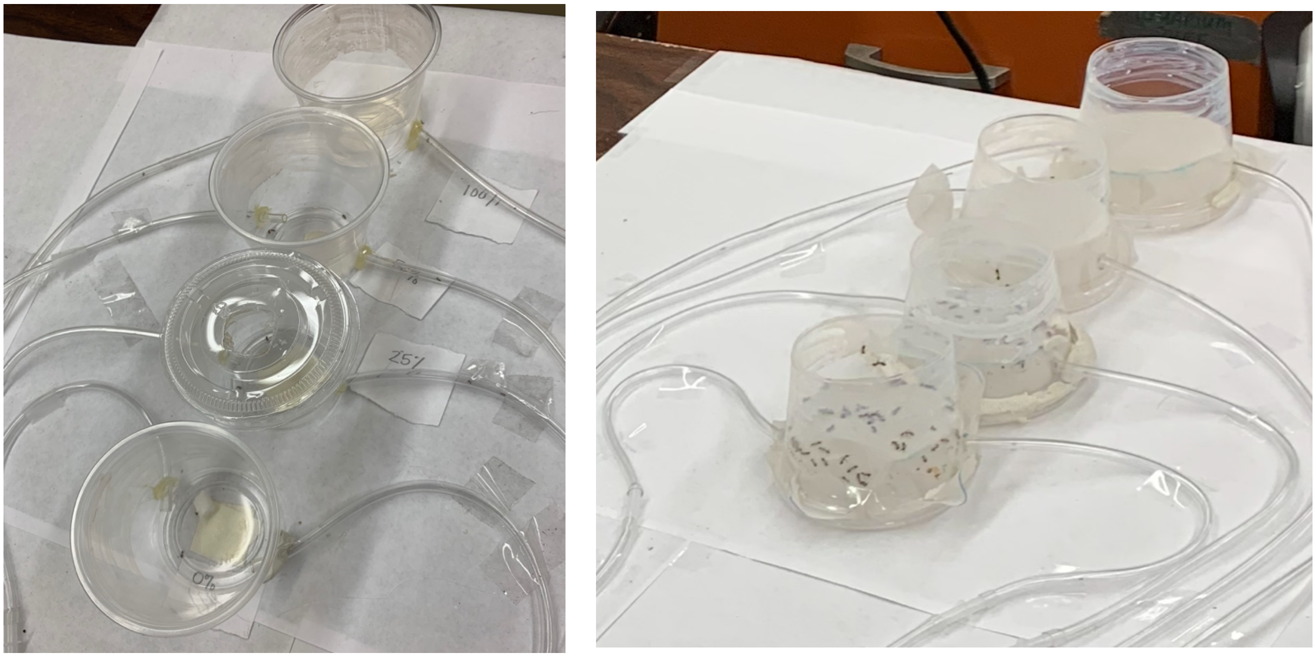


**Fig. S2:** Picture of cells where stimuli were placed along experimental grids. Left represents trials where formic acid was used; highlighting inside of the cell where formic acid was placed as well as the lidding of cells to prevent formic acid from dissipating over time. Right represents trials where *L. occidentale* was used. Mesh prevented *L. humile* and *L. occidentale* from coming into physical contact, but pheromones could be exchanged.
